# Common neural bases for processing speech prosody and music: An integrated model

**DOI:** 10.1101/2021.05.12.443804

**Authors:** Alice Mado Proverbio, Elisabetta Piotti

## Abstract

It is shared notion that speech and music processing share some commonalities. Brain bioelectrical activity was recorded in healthy participants listening to music obtained by digitally transforming real speech into melodies played by viola. Sentences were originally pronounced with a positive or negative affective prosody. The research’s aim was to investigate if the emotional content of music was extracted similarly to how the affective prosody of speech is processed.

EEG was recorded from 128 electrodes in 20 healthy students. Participants had to detect rare neutral piano sounds while ignoring viola melodies. Stimulus negative valence increased the amplitude of frontal P300 and N400 ERP components while a late inferior frontal positivity was enhanced in response to positive melodies. Similar ERP markers were previously found for processing positive and negative music, vocalizations and speech. Source reconstruction applied to N400 showed that negative melodies engaged the right superior temporal gyrus and right anterior cingulate cortex, while positive melodies engaged the left middle and inferior temporal gyrus and the inferior frontal cortex. An integrated model is proposed depicting a possible common circuit for processing the emotional content of music, vocalizations and speech, which might explain some universal and relatively innate brain reaction to music.

## INTRODUCTION

Several studies have shown that music and speech processing rely on shared brain resources, for example for the decoding of their temporal (Abrams et al., 2011) or syntactical structure (Koelsch et al., 2013; Kunert et al., 2015; Steinbeis and Koelsch, 2008; Sun et al., 2018). However, the issue of whether emotion processing in speech and music is similarly grounded remains rather unexplored. Some authors have argued that the reason the emotional content of specific music pieces can be recognized by any listener, regardless of their age, culture and education (Dalla Bella et al., 2001; Flom et al., 2008; Fritz et al., 2009) therefore in a rather universal and innate manner, relies on this commonality (e.g., Proverbio et al., 2020a). Panksepp and Bernatzky (2002) hypothesized that music ability reflects the ancestral tendency of the mammalian brain to transmit and receive basic emotional sounds. For example, it seems that minor chords share the same harmonic relationships that are found in sad speech prosody (Cook et al., 2006). At this regard, Paquette and coworkers (2018) proposed that music could tap into neuronal circuits that have evolved primarily for the processing of biologically important emotional vocalizations. A few studies have indeed found common brain activations (including middle and superior temporal cortex, medial frontal cortex and cingulate cortex) during listening to both non-linguistic vocalizations and musical excerpts (Aubé et al., 2015; Escoffier et al., 2013; Belin et al., 2008; Sachs et al., 2018, Paquette et al., 2018). Although from these studies it can be clearly inferred that similar mechanisms support the comprehension of the emotional content of vocalizations and music (Paquette et al., 2020), there is not a precise indication about specific similarities/differences in neural processing and its time course, because of the different structure of stimuli usually considered. Moreover, while many ERP studies investigated the brain ability to detect a mismatch between happy vs. sad/angry prosodic information (e.g., Besson et al., 2002; Bostanov et al., 2004; Paulmann and Kotz, 2008), in our knowledge, not many studies have provided clear ERP markers of negative vs. positive emotional connotation of auditory information (Schapkinet al., 2000; Schmidt and Trainor, 2001).

This study is part a broader research project designed to compare the neural processing of speech (meaningful sentences), instrumental string music and affective vocalizations (e.g., laughter and crying), characterized by a positive vs. negative affective valence, with the aim of understanding how the brain extracts the affective meaning of auditory information.

In the present study, we investigated this issue by measuring brain potentials in response to music derived from speech, originally positive and negative, both in semantics and prosody. The ERP data recorded to speech stimuli are reported in Proverbio et al.’s (2020b) paper. Indeed speech (besides meanings and semantics) also conveys emotional information through the modulation of the tone of voice, the speech intonation and speed, and the affective prosody (Nygaard and Queen, 2008; Kotz and Paulmann, 2007), the so called “melody of language” (Wildgruber et al., 2005; Brück et al., 2011).

Prosody is characterized by variations in the suprasegmental characteristics of language, such as tones, the duration of syllables and the quality of the voice (Banse and Scherer, 1996). The neural circuits involved in the processing of affective prosodic information includes the superior temporal cortex (Wildgruber et al., 2005; Beacousin et al., 2006; Früholtz and Grandjean., 2013; Witteman et al., 2012; Kotz et al., 2013), particularly over the right hemisphere (Zhang et al., 2018; Kotz et al., 2003; Wildgruber et al., 2005; Kotz et al., 2013), and the right prefrontal cortex (Wildgruber et al., 2005; Ethofer et al., 2006; Brück et al., 2011, Johnstone et al., 2006). Overall, the processing and comprehension of affective prosody seem to engage mostly right than left hemispheric structures (Schirmer and Kotz, 2006, Wildgruber et al., 2005; Beacousin et al., 2006; Ethofer et al., 2006; Van Lancker e Sidtis, 1992). For example, Kotz and coworkers (2003) recorded fMRI signals in subjects listening to semantically neutral sentences pronounced with neutral, positive or negative intonation. The activation data showed a left temporal and left inferior frontal activation for the neutral intonation, and bilateral activation (engaging the homologous right hemispheric areas) for the processing of affective prosody (Kotz et al., 2003).

In addition to the prefrontal/superior temporal circuits, both clinical (Paulmann et al., 2010) and neuroimaging studies have suggested the involvement of the orbitofrontal cortex (OFC) (Brück et al., 2011; Witteman et al., 2012; Wildgruber et al., 2005; Ethofer et al., 2006) and the inferior frontal cortex (Zhang et al., 2018) in prosody comprehension. Limbic structures such as the amygdala (Sander et al., 2005; Beacousin et al., 2006) also play a crucial role.

As for the neural mechanisms subtending the ability to comprehend the affective content of music, positive music seems to engage the inferior frontal gyrus, the superior anterior insula and the ventral striatum (Koelsch et al., 2006). In the EEG/fMRI study by Flores-Gutiérrez and collaborators (2007) a hemispheric asymmetry was found for the processing of pleasant and unpleasant music with pleasant music activating the left posterior temporal-parietal region, and the left medial prefrontal area, and the unpleasant music activating right superior frontopolar region, the insula (slightly asymmetrically) and the right auditory area. Similarly Khalfa and coauthors (2008) examined patients with right or left anterior medial temporal resections finding that while patients with left lesions showed a significant deterioration in the identification of happy musical pieces patients with resection of the right anteromedial temporal lobe showed deficits in the recognition of sadness. The authors concluded that the right anteromedial temporal region was involved in the recognition of negative emotions, while the left anteromedial temporal region processed both positive and negative emotions (Khalfa et al., 2008).

The aim of the present study was to record the brain bioelectric responses of subjects listening to melodies extracted from verbal messages, with positive or negative value, in order to identify the neural mechanisms supporting the processing of affective music. The hypothesis was that, if music affective content was processed similarly to how the emotional content of vocalizations and speech are, this would result in similar ERP markers. For example, N400 and Late Positivity components of ERPs were found larger in response to negative (N400) and positive (LP) vocalizations, music and speech stimuli in previous electrophysiological investigations (e.g. Proverbio et al., 2020a; Proverbio et al., 2020b). Again, in different paradigms, Chen and coworkers (2008) found larger negativities (500-700 ms) associated to the listening of negatives vs. positive music. Speculalry, Schapkin and coworkers (2000) found larger P2 and P3 positive peaks to positive than to negative words, which is compatible with the aforementioned pattern of electrophysiological results. Besides enhanced N400 and LP potentials in response to emotions of opposite valence, we expected to find possible evidences a right hemispheric asymmetry for the processing of negative music, and a left hemispheric asymmetry for the processing of positive music, as predicted by the available literature (e.g., Altenmüller, et al., 2002; Khalfa et al., 2008; Koelsch et al., 2006).

In this study, a source reconstruction was also performed on the electrophysiological data, in order to investigate the inner brain sources involved in the processing of positive and negative musical stimuli. The results of the inverse solution were compared across studies, and with the findings obtained by previous fMRI investigations. This allowed the creation of a comprehensive model of how emotional nuances of auditory information are extracted and comprehended, regardless of the specific sound nature (speech, vocalizations and music).

## MATERIAL AND METHODS

### Participants

Twenty university students (10 men and 10 women) ranging in age from 18 to 35 years (mean age= 23.45; SD= 2.3) took part in the study. All participants had normal or corrected-to-normal vision and normal hearing. They were strictly right-handed as assessed by the Oldfield Inventory (mean score= 0.73; SD= 0.17. Max= +1, Min= -1) and reported no history of neurological or mental disorders. Experiments were conducted with the understanding and written consent of each participant according to the Declaration of Helsinki (BMJ 1991; 302: 1194), with approval from the Ethical Committee of the local University (RM-2019-176). The data from 3 participants were subsequently discarded from ERP analyses for excessive EEG artifacts. The final sample of 17 subjects had a mean age of 23.65 (SD= 2.3).

### Stimuli

To create the music stimuli, the speech stimuli used in the study by Proverbio et al. (2020b) were digitally converted into instrumental music for viola. The original stimuli were 223 sentences with positive (n= 99), negative (n= 99) and neutral (n= 25) emotional content, spoken by two professional male and female speakers. In Table 1 are listed some of the sentences used in this neurolinguistics study.

**Table 1.**
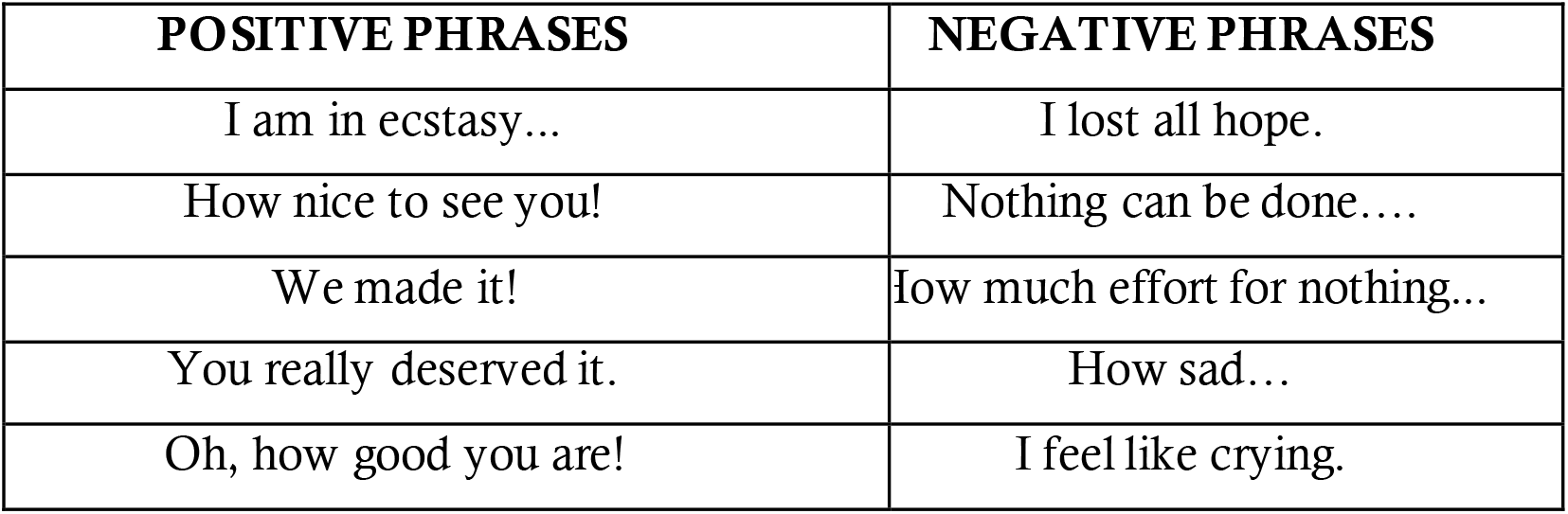
Examples of spoken sentences (taken from Proverbio et al., 2020b study) from which prosodic information was extracted and digitally converted into viola played melodies, as a function of their emotional content.

Original verbal utterances included expressions of euphoria, surprise, encouragement, gratitude, appreciation, hope and relief for the positive category, and self-pity, boredom, frustration, depression, despair, discouragement and reaction to physical pain for the negative category. Two professional speakers recorded stimulus sentences. They were instructed to read the text spontaneously and with participation, by identifying with the characters involved in depicted scene (at the Stanislavskij manner). A microphone (*Tonor brand*) and audio file editor *Audacity* were used to record audio tracks. The files were saved in WAV format. The two stimulus categories, positive and negative, were matched for number of letters and words, as demonstrated by two 1-way ANOVAs carried out across categories (number of words: [F (1,98)= 0.71795, p= 0.39; number of letters [F(1,98)= 1.3923, p= 0.24]). The average number of words per sentence was 3.28, while the average number of letters per sentence was 15.89, with no difference between the positive and negative categories.

Audio tracks lasted exactly 2200 ms, were normalized, and levelled (frequency of 44100 Hz, resolution of 16 bits). Their acoustic intensity was measured through a PCE-999 sound level meter. Values were 59.2 dB (SD= 3.8) for positive utterances and 56.5 dB (SD= 4.3) for negative utterances. Intensity values did not statistically differ across classes, as proofed by a 1-way ANOVA (p= 0.99).

*Apple’s Logic Pro X* program, a professional software for producing and recording music, was used to transform speech stimuli into musical stimuli. The individual stimuli were imported into the program and, subsequently, using the “Turn on Flex” function and selecting the “Flex Pitch” option, each stimulus was converted into a MIDI track (“Create MIDI from flex data”). As for the timbre of the track, the sound of the strings “King’s Cross”, from the “Studio Strings” section (which roughly corresponds to viola timbre) was used for negative and positive stimuli. For neutral stimuli, which were then used as task-related target stimuli, the “Steinway Grand Piano” piano tone was selected. Furthermore, all the stimuli were transposed 12 semitones higher than the original melody of the sentences, to increase the similarity with the viola instrument. Each stimulus lasted 2.20 seconds, the same as the original stimuli, with a resolution of 16 bits.

To make the end of each stimulus more “natural”, *Audacity* program was used to attenuate the last 150 ms of the melody, through the “Fade out” function.

The same procedure was used on other 50 stimuli with the same characteristics (positive, n= 20; negative, n= 20; neutral, n= 10), to be used in the training phase.

### Stimulus validation

The stimuli were preliminarily evaluated for their valence and their arousal. Fifteen judges (8 males and 7 females), aging between 22 and 28 years (mean= 24.06; SD= 1.95), non-musicians, listened to the 198 positive and negative musical stimuli (extracted from speech utterances as detailed above). Each judge was asked to indicate, the perceived emotional value of each melodic fragment heard, by means of a 3 points Likert scale (0= negative valence; 1= neutral valence; 2 = positive valence). They were also asked to indicate how much they felt activated by the stimulus, using a modified version of the Manikin test by Bradley and Lang (1994) (0 = calm; 1 = activated; 2 = very activated).

Two separate two-ways ANOVAs were performed on the obtained valence and arousal scores, as a function of stimulus type. In the valance analysis the factors of variability were emotion (2 levels: positive, negative) and valence (3 levels: negative, neutral, positive). The results of the analysis showed a significant interaction between the two factors [F (2,392) = 4.699, p< 0.005]. The HSD Tukey test showed that, although both negative and positive stimuli were more frequently judged as neutral than affective, negative stimuli were classified as negative more frequently than positive stimuli and that positive stimuli were classified as positive more frequently than the negative stimuli. The ANOVA performed on the arousal levels examined two factors of variability: emotion (2 levels: positive, negative) and arousal (3 levels: calm, activated, very activated). The ANOVA yielded the interaction of emotion x arousal [F (2, 392) = 29.026, p< 0.000]. Tukey post-hoc comparisons showed that negative stimuli were classified as activators more frequently than positive stimuli (neg-activ= 8.18, SD= 0.25; pos-activ= 6.65, SD= 0.25). Finally, positive stimuli were classified as very activating more frequently than the negative stimuli (neg-very activ. = 3.35, SD = 0.32; pos-very activ.= 6.19, SD= 0.32).

### Stimulus material and procedure

The 223 musical clips (99 negative, 99 positive, and 25 neutral) were binaurally presented in random order within six experimental runs. Some examples of auditory stimuli are given in the Supplementary files. Each run comprised 37 stimuli, except for the first one, which comprised 38 stimuli. In each run, half stimuli were positive and the other half were negative in valence. They were randomly intermixed with min 3/max 5 targets (for each run). The ISI (Inter Stimulus Interval) was 600 ms, with a random interval of 200 ms (600 ± 200 ms).

EEG recordings were performed in an electrically shielded anechoic chamber (a Faraday cage). Stimuli presentation and triggering was performed using *Eevoke* Software for audiovisual presentation (*ANT Software*, Enschede, The Netherlands). Participants listened to the stimuli through a set of *Sennheiser HD202* headphones. While listening to auditory stimuli participants observed a visual background projected on a PC screen (a colored image of a living room); this expedient was used to prevent excessive alpha waves from being generated over the occipito/parietal area. The participants were instructed to fixate the center of the screen located about 114 cm from their eyes, to relax, not to contract face or body muscles and to avoid blinking as much as possible in order to get a good EEG signal.

The participant’s task was to press a key with their index finger, as accurately and quickly as possible, whenever they heard the sound of a piano, while non-targets stimuli were viola melodies. The left and right responding hands were used alternately throughout the recording session, and the order of the hand and task conditions were counterbalanced across subjects. Prior EEG recording sessions participants underwent a training session in which they familiarized with experimental setting.

### EEG recordings and analysis

The EEG was recorded and analyzed using *EEProbe* recording software (*ANT Software*, Enschede, The Netherlands). EEG data were continuously recorded from 128 scalp sites according to the 10–5 International System. Sampling rate was 512 Hz. Horizontal and vertical eye movements were additionally recorded, and linked ears served as the reference lead. Vertical eye movements were recorded using two electrodes placed below and above the right eye, while horizontal movements were recorded using electrodes placed at the outer canthi of the eyes, via a bipolar montage. The EEG and electro-oculogram (EOG) were filtered with a half-amplitude band pass of 0.016–70 Hz. Electrode impedance was maintained below 5 KOhm. EEG epochs were synchronized with stimulus onset and analyzed using *ANT-EEProbe software*. Computerized artifact rejection was performed prior to averaging to discard epochs in which amplifier blocking, eye movements, blinks or excessive muscle potentials occurred. The artifact rejection criterion was a peak-to-peak amplitude exceeding 50 μV and resulted in a rejection rate of ∼5%. Event-related potentials (ERPs) were averaged offline from 100 ms before to 2000 ms after stimulus onset. ERP components were identified and measured with respect to the average baseline voltage over the interval from −100 to 0 ms.

Isocolour topographic maps of the surface voltage of main ERP components were computed. A LORETA (Low Resolution Electromagnetic Tomography, Pasqual-Marqui et al., 1994), was also applied to the ERP responses obtained in the time window between 1100 and 1200 ms. An advanced version of standardized LORETA was used, the swLORETA (Palmero-Soler et al., 2007), with the following properties of the reference space: grid spacing (grid spacing)= 5 mm; Tikhonov regularization: signal-to-noise ratio= 3. LORETA is a discrete linear solution to the inverse EEG problem, and it corresponds to the 3D distribution of neural electric activity that maximizes similarity (i.e. maximizes synchronization) in terms of orientation and strength between neighboring neuronal populations (represented by adjacent voxels). In this study, an improved version of standardized weighted low-resolution brain electromagnetic tomography was used; this version incorporates a singular value decomposition-based lead field weighting method (i.e. swLORETA, Palmero-Soler *et al*., 2007). The source space properties included a grid spacing (the distance between two calculation points) of 5 points and an estimated signal-to-noise ratio of 3, which defines the regularization (higher values indicating less regularization and therefore less blurred results). The use of a value of three to four for the computation of SNR in the Tikhonov’s regularization (1963) produces superior accuracy of the solutions for any inverse problems assessed. swLORETA was performed on the group data (grand-averaged data) to identify statistically significant electromagnetic dipoles (p< 0.05) in which larger magnitudes correlated with more significant activation. The data were automatically re-referenced to the average reference as part of the LORETA analysis. A realistic boundary element model (BEM) was derived from a T1 -weighted 3D MRI dataset through segmentation of the brain tissue. This BEM model consisted of one homogeneous compartment comprising 3446 vertices and 6888 triangles. Advanced source analysis (ASA) employs a realistic head model of three layers (scalp, skull and brain) created using the BEM. This realistic head model comprises a set of irregularly shaped boundaries and the conductivity values for the compartments between them. Each boundary is approximated by a number of points, which are interconnected by plane triangles. The triangulation leads to a more or less evenly distributed mesh of triangles as a function of the chosen grid value. A smaller value for the grid spacing results in finer meshes and vice versa. With the aforementioned realistic head model of three layers, the segmentation is assumed to include current generators of brain volume, including both gray and white matter. Scalp, skull, and brain region conductivities were assumed to be 0.33, 0.0042 and 0.33, respectively (Zanow and Knösche, 2004). The source reconstruction solutions were projected onto the 3D MRI of the Collins brain, which was provided by the Montreal Neurological Institute. The probabilities of source activation based on Fisher’s *F*-test were provided for each independent EEG source, the values of which are indicated in a so-called ‘unit’ scale (the larger, the more significant). Both the segmentation and generation of the head model were performed using the ASA software program (*Advanced Neuro Technology, ANT*).

The mean area amplitude of P2 response was quantified at F5-F6, F7-F8 and AF3-AF4 electrode sites in between 200 and 300 ms. The mean area amplitude of P300 response was quantified at AF7-AF8 and F3-F4 electrode sites in between 600 and 700 ms. The mean area amplitude of N400 response was quantified at AFF1-AFF2 and FFC1h-FFC2h electrode sites in between 1100 and 1200 ms. The mean area amplitude of Late Positivity was quantified at AFz, Fz and AFF1-AFF2 electrode sites in between 1700 and 1800 ms. ERP components were measured when and where they reached their maximum amplitude at scalp sites, and according to previous literature on emotional prosody and music.

Separate multifactorial repeated-measure analyses of variance (ANOVAs) were performed on the values computed in the various time windows. The factors were “valence” (positive, negative), “electrode” (dependent on the ERP component of interest) and “hemisphere” (left hemisphere, or LH; right hemisphere, or RH).

The average reaction time to target stimuli was 1052.4 ms; the errors detected were minimal, with an average accuracy of 98.2%.

## RESULTS

The ANOVA carried out on P2 (200 – 300 ms) amplitude values did not show any significant effect of valence, thus indicating the negative and positive melodies were perceptually matched and virtually similar, at the first stages of auditory processing.

The ANOVA evidenced a significant main effect of valence on the amplitude of the P300 (600–700 ms) component [F (1,16)= 4.65; p<0.047], showing larger amplitude in response to negative (2.03 μV, SD= 0.45) than positive melodies (1.50 μV, SD = 0.45) as visible in Fig. 1. The ANOVA also yielded the significance of hemisphere factor [F (1,16)= 6.13; p<0.024], with greater P300 responses over right (2.11 μV, SD = 0.54) than left (1.41 μV, SD = 0.35) scalp areas.

**Figure 1.**
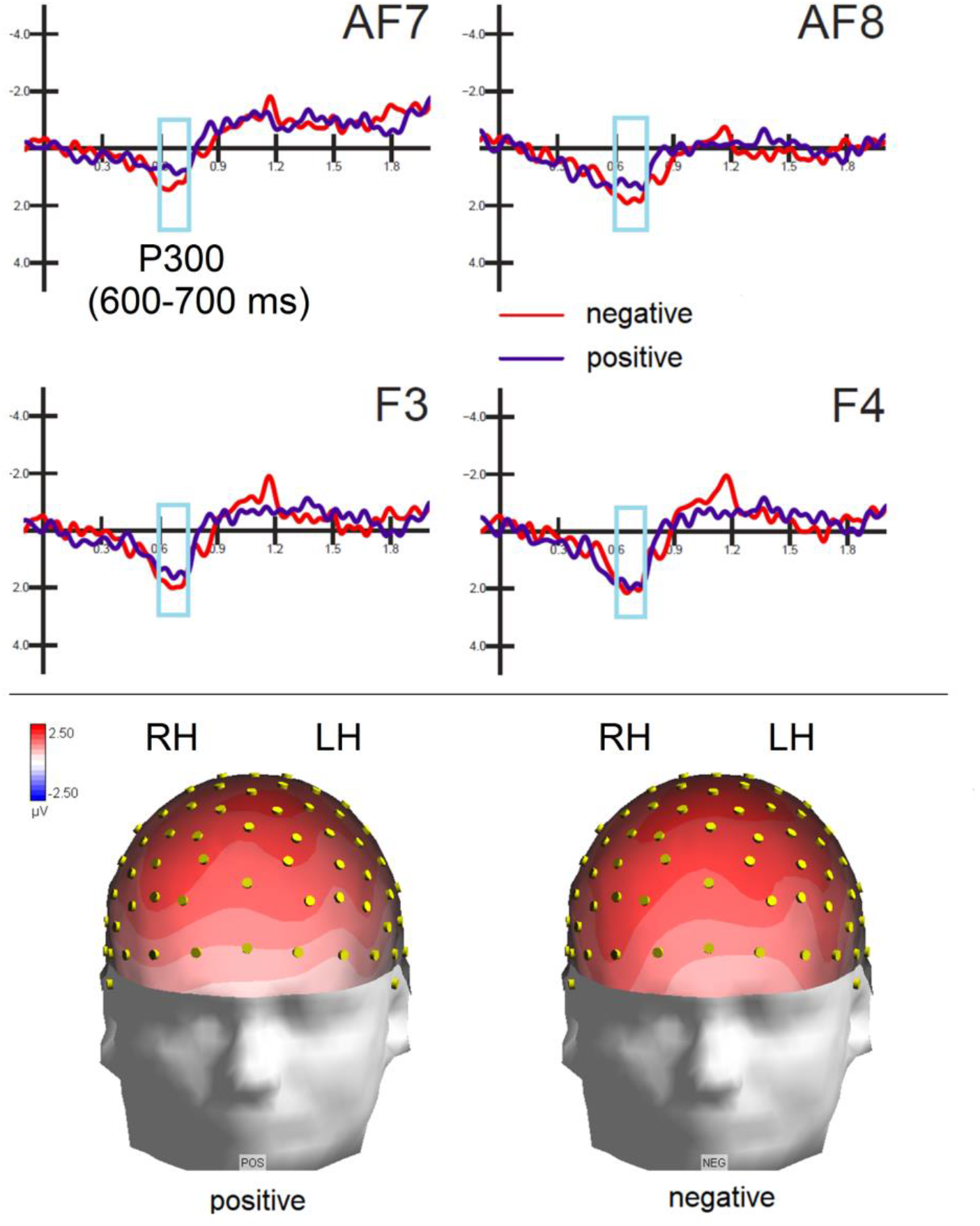
(Top) Grand-average ERP waveforms elicited by negative and positive melodies as recorded at left and right anterior frontal and frontal sites. (Bottom) Isocolour topographical maps (front view) of P3 response at peak latency (670 ms). It can be appreciated the marked right hemispheric asymmetry.

The ANOVA carried out on N400 (1100–1200 ms) evidenced a significant main effect of valence [F (1,16)= 6.40; p<0.022] with greater amplitudes to negative (−2.04 μV, SD = 0.44) than positive melodies (−0.86 μV, SD= 0.42) as can be appreciated in Fig. 2. The further significance of Electrode [F (1,16)= 14.30; p<0.001] showed that N400 was larger over anterior than frontal sites (AFF1-AFF2= -1.65 μV, SD= 0.39; FFC1h-FFC2h= -1.25 μV, SD= 0.34).

**Figure 2.**
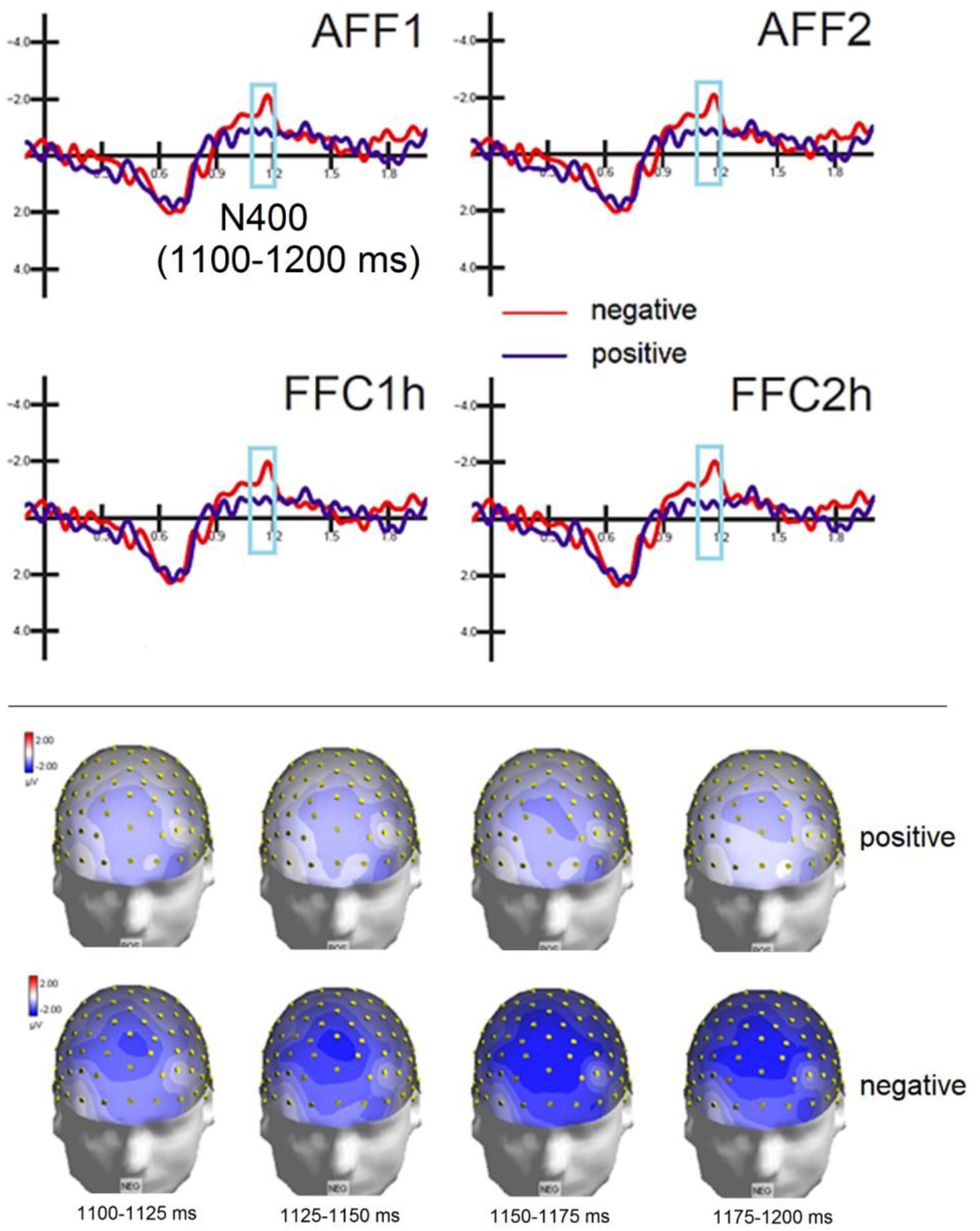
(Top) Grand-average ERP waveforms elicited by negative and positive melodies as recorded at left and right anterior frontal and frontocentral sites. (Bottom) Time series of isocolour topographical maps (front view) of N400 voltage recorded to positive and negative melodies in between 1100 to 1200 ms post-stimulus.

The ANOVA carried out on *Late Positivity* (LP, 1700-1800 ms) evidenced a significant main effect of valence [F (1,16) = 6.60; p<0.02] with greater amplitudes to positive (−0.73 μV, SD= 0.59) than negative melodies (−1.55 μV, SD= 0.54), as can be appreciated by looking at ERP waveforms and topographical maps of Fig 3.

**Figure 3.**
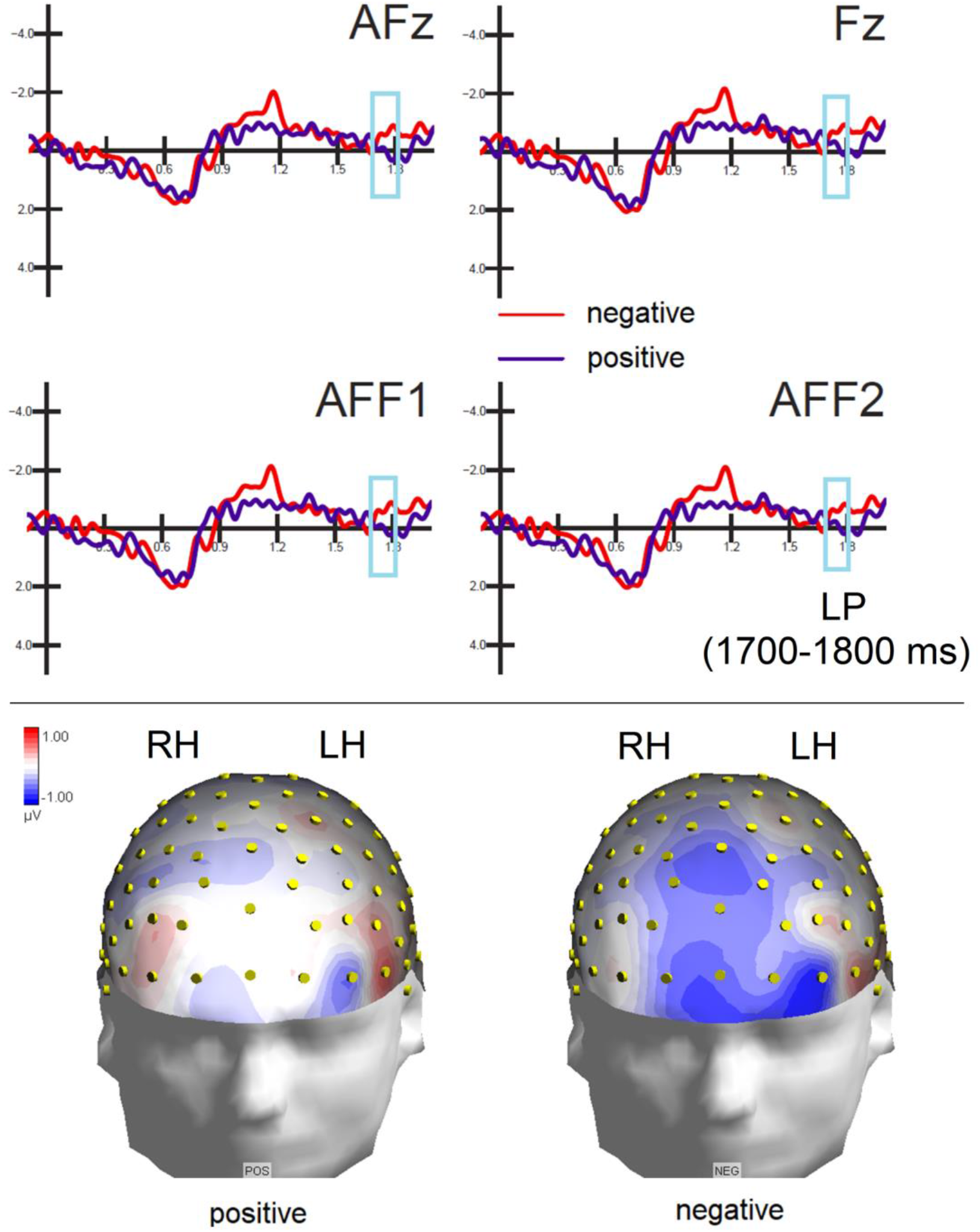
(Top) Grand-average ERP waveforms elicited by negative and positive melodies as recorded at midline anterior frontal and frontal sites, and left and right anterior frontal sites. (Bottom) Isocolour topographical maps (front view) of LP amplitude measured at peak latency (1745 ms).

To identify the neural generators of the intracranial activity associated with valence processing at N400 level, two swLORETAs (*standardized weighted Low Resolution Electromagnetic Tomography*) were performed on the brain responses recorded between 1100 e 1200 ms in response to positive and negative melodies. This time latency was selected because it showed the largest effect of emotional valence, in terms of amplitude modulation, but of course was not sensory specific, which may explain why Heschl’s gyri (BA41) were found not to be particularly active at this later stage. Table 2 shows the active electromagnetic dipoles explaining the surface voltage while source reconstruction is displayed in Fig. 4.

**Table 2.** Talairach coordinates (in mm) of electromagnetic dipoles corresponding to the intracranial generators explaining the surface voltage recorded during the 1100–1200 ms time window in response to positive (Top) and negative (bottom) melodies.

| NEGATIVE |  |  |  |  |  |  |  |  |
| --- | --- | --- | --- | --- | --- | --- | --- | --- |
| Magn. | T-x<br>[mm] | T-y<br>[mm] | T-z<br>[mm] | Hem. | Lobe | Gyrus | BA | Presumed function |
| 11 .29 | -38 .5 | 46 .3 | -2 .3 | L | F | Middle Frontal | 10 | Attention and working memory<br>ENCODING |
| 11.26 | -38 .5 | 43 .4 | 23 .9 | L | F | Middle Frontal | 10 |  |
| 9 .88 | -58 .5 | 3 .3 | 20 .5 | L | F | Precentral | 6 |  |
| 9 .86 | -58 .5 | -39 .6 | 25 .1 | L | P | Inferior Parietal lobule | 40 | Affective or dissonant music processing (Minati et al., 2009; Bogert et al., 2016) |
| 9 .35 | 40 .9 | 55 .3 | 7 | R | F | Middle Frontal | 10 |  |
| 8 .78 | 40 .9 | 43 .4 | 23 .9 | R | F | Middle Frontal | 10 |  |
| 8 .38 | 60 .6 | -28 .5 | 17 .1 | R | T | Superior Temporal | 42 | MUSIC<br>(Negative valence) |
| 6 .46 | 11 .3 | 35 .3 | 5 .3 | R | Limbic | Anterior Cingulate | 24 |  |
| 6 .18 | 60 .6 | -58 .9 | 14 .5 | R | T | Superior Temporal | 22 |  |
| 6 .07 | -38 .5 | -86 .4 | -12 .4 | L | O | Inferior Occipital | 18 |  |
| 6 .00 | -28 .5 | -82 .1 | 39 .5 | L | P | Precuneus | 19 |  |
| 5 .00 | 21 .2 | -0 .6 | -28 .2 | R | Limbic | Uncus | 36 | Emotion |
| POSITIVE |  |  |  |  |  |  |  |  |
| Magn. | T-x<br>[mm] | T-y<br>[mm] | T-z<br>[mm] | Hem. | Lobe |  | BA | Presumed function |
| 1 .09E-09 | -48 .5 | 45 .3 | 6 .1 | L | F | Middle Frontal | 46 | Attention and working memory<br>ENCODING |
| 8 .84 | -38 .5 | 43 .4 | 23 .9 | L | F | Middle Frontal | 10 |  |
| 6 .48 | -58 .5 | -24 .5 | -15 .5 | L | T | Middle temporal | 21 | MUSIC<br>(Positive valence) |
| 5 .86 | -58 .5 | -8 .7 | -21 .5 | L | T | Inferior Temporal | 20 |  |
| 5 .61 | -38 .5 | -15 .3 | -29 .6 | L | T | Inferior Temporal | 20 |  |
| 5 .93 | 11 .3 | 30 .5 | 49 .8 | R | F | Superior Frontal | 6 |  |
| 5 .48 | 60 .6 | -13 | 27 .7 | R | P | Postcentral | 3 |  |
| 5 .12 | 50 .8 | 45 .3 | 6 .1 | R | F | Middle Frontal | 46 |  |
| 5 .06 | 60 .6 | 14 .3 | 12 .5 | R | F | Inferior Frontal | 44 | Positive vocalizations (Proverbio et al., 2020a; Warren et al., 2006; Belin et al., 2008), Prosody (Kotz et al., 2003), Pleasant music (Koelsch et al., 2006) |
| 5 .02 | 21 .2 | -8 | -28 .9 | R | Limbic | Uncus | 36 | Emotion |

**Figure. 4.**
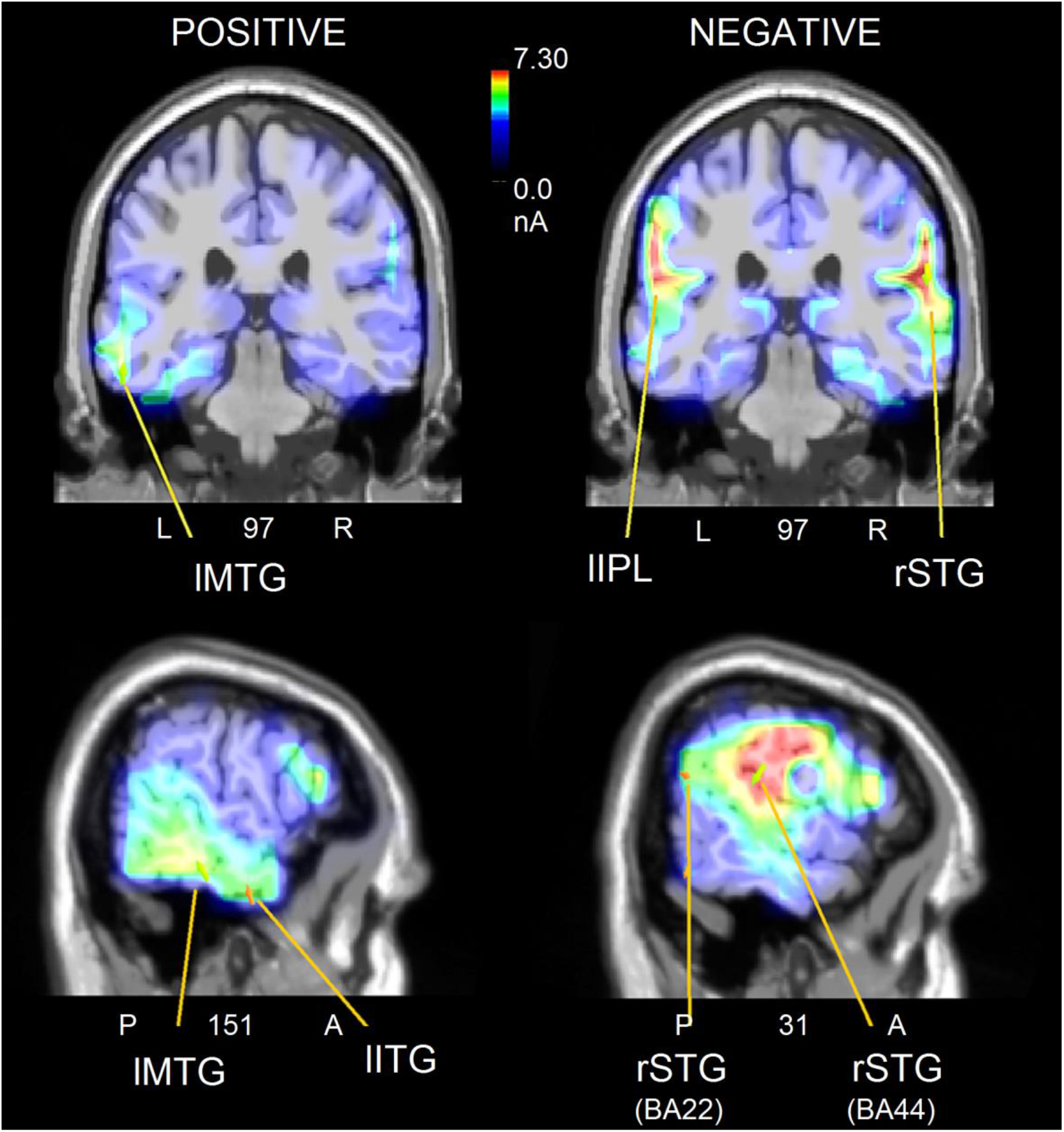
Coronal (top) and sagittal (bottom) views of active sources during the processing of positive and negative musical excerpts in between 1100 to 1200 ms latency post -stimulus. The different colors represent differences in the magnitude of the electromagnetic signal (in nAm). The electromagnetic dipoles are shown as arrows and indicate the position, orientation and magnitude of the dipole modeling solution applied to the ERP waveform within the specific time window. The numbers refer to the displayed brain slice in sagittal view: A= anterior, P= posterior. The images highlight the strong activation of the left MTG for positive music and of the right STG for negative music.

For both conditions, the left middle frontal gyrus was activated, possibly indicating attentional allocation and working memory processing. As for the elaboration of the emotion underlying the melody, negative melodies mainly engaged the right superior temporal gyrus and the right cingulate cortex, while the positive melodies mainly engaged the left middle temporal and inferior temporal gyri.

## DISCUSSION

The goal of the research was to investigate the neural mechanisms involved in the extraction of the emotional meaning of musical information and to identify the neural circuits involved in the processing of musical stimuli with different emotional valence, possibly associated with the homologous mechanism of prosody comprehension. At this aim, negative and positive spontaneous speech utterances were digitally converted into viola melodies thus maintaining the same melodic profile, but being strictly musical in nature. Stimulus validation revealed that listeners were not aware of the emotional content of musical melodies, since they judged the stimuli as neutral more frequently than affective. Indeed, to an unaware listening, they seem fragments of Paul Hindemith’s viola music, slightly dissonant. However, negative stimuli were judged as negative more frequently than positive stimuli and vice versa. This finding is intriguingly similar to what found by Belin and coworkers (2008) who analyzed fMRI signals of individuals who listened to animal (monkeys and cats) and human vocalizations recorded in positive or in negative contexts. The authors found that although the study participants were not able to explicitly distinguish the “positive” and “negative” affective vocalizations of animals, the two types of stimuli were clearly distinguished in their brain (Belin et al ., 2008). fMRI data revealed that the emotional valence modulated the activation of the secondary auditory cortex, with greater signals for negative than positive animal vocalizations. The same pattern of results was found for the processing of human vocalizations, in the same study. They hypothesized that this could depend on the fact that negative vocalizations were characterized by longer sounds that activated the auditory cortex more effectively. Also in our study, although not fully comprehended at a conscious level by the listeners, the intrinsic negative and positive nature of music melodies strongly affected brain potentials. The analysis of ERP amplitudes recorded between 200 and 300 ms did not show any significant effect of valence, thus indicating that two types of melodies were virtually similar and matched from the perceptual point of view. On the other hand, later anterior P300 response was of greater amplitude to negative than positive stimuli. This component might reflect the attentional orienting toward the stimuli (Proverbio et al., 2016; Putkinen et al., 2017). Since viola melodies were non-targets and had to be ignored, this finding might indicate that negative stimuli were able to attract attention to a greater extent than positive stimuli. Regardless of valence P300 was larger over the right hemisphere, which might be also interpreted as a general right hemispheric dominance for processing music melodies in non-musicians (Samson and Zatorre, 1988; Zatorre et al., 2002: Proverbio et al., 2016). Interestingly the same hemispheric asymmetry was found for speech prosody and the processing of intonation and language inflection (Ross, 1981; Ross et al., 1997; Wildgruber et al., 2005; Ethofer et al., 2006; Zhang et al., 2018).

The second component investigated was the N400, recorded between 1100 and 1200 ms, which was influenced by stimulus valence: the negative melodies elicited a much larger N400 response than the positive melodies. This result is highly consistent with what found by Proverbio et al., (2020a) comparing positive (e.g., laughs) and negative vocalizations (e.g., crying), and Proverbio et al. (2020b) comparing positive (<<How nice to see you!>>) and negative speech utterances (e.g.: <<How sad…>>. In both studies, stimuli with a negative valence generated larger N400 responses than stimuli with a positive valence. In this study, despite the lack of the linguistic content, the melodic profile of musical fragments possibly reflected the prosodic structure of the sentences from which they were derived, which might explain the data similarities. This hints at a common neural mechanism for extracting affective information from vocalizations, speech and music. It has indeed been proposed that vocalizations and music use similar acoustic patterns to express emotional information (Juslin and Laukka, 2003)

Finally, the Late Positivity (LP) recorded between 1700 and 1800 ms was larger to positive than negative melodies, which, again, is strikingly similarly to what found for nonverbal vocalizations as well as speech utterances by previous ERP investigations (Proverbio et al., 2020a; Proverbio et al., 2020b). The inferior frontal distribution of this component (larger to positive stimuli) also agrees with the fMRI findings by Belin and coauthors (2008) for human and animal vocalizations and Koelsch and coauthors (2006) contrasting unpleasant and pleasant music processing, and with source reconstruction data reported below.

Since the valence-dependent amplitude modulation was greater at N400 level a source reconstruction was applied to brain potentials recorded at this latency stage (1100 and 1200 ms) to unravel the neural circuits subtending the understanding of the emotional content of music. For both listening conditions, the most active dipole was the left middle frontal gyrus (MFG, BA 10). In several neuroimaging studies (e.g., Altenmüller et al., 2014; Proverbio and De Benedetto, 2018; Bogert et al., 2016) it was found a large MFG activation during listening to music with a strong emotional content. This activation could reflect coding processes and attention to the stimuli, linked to working memory. Also, the left hemispheric asymmetry in prefrontal activation is compatible with the HERA model by Tulving (1994) who hypothesized a role of the left prefrontal area in the encoding of incoming information (as opposed to memory retrieval) (see also Green et al., 2012). These findings are also compatible with the model proposed by Heller (1993), and the EEG data by Rogenmoser and coauthors (2016) arguing that the frontal lobe is involved in modulating valenced experiences, with a the left hemispheric asymmetry for positive emotions.

Another region active in both listening condition was the right uncus (BA 36), a limbic region that is probably involved in the processing of emotions and memory (Catani et al., 2013). According to the literature, the uncus is also quite active during listening to vocalizations (Vassal et al., 2016; Proverbio et al., 2020a).

Besides frontal areas, the most active source during listening to negative melodies was the left inferior parietal lobule (BA 40), which is thought to be engaged in the processing of speech utterances (Friederici, 2011). This area has also been found active in the study by Proverbio et al. (2020b), in which participants listened to speech with a negative content, as well as in Minati et al.’s study (2009), in which participants listened to dissonant music; finally, in participants listening to affective music (Bogert et al., 2016).

Overall, the source reconstruction data provided evidence of a hemispheric asymmetry for the processing of positive vs. negative melodies (similarly to what found, for example, by Altenmüller, et al., 2002). Overall, listening to positive melodies engaged the left middle temporal gyrus (BA 21) and left inferior temporal gyrus (BA 20); on the contrary, music with negative valence engaged more the right superior temporal gyrus (BA 42 and BA 22) and the right anterior cingulate cortex (BA 24). This pattern of lateralization is consistent with the hypothesis proposed by various authors according to which positive emotions (and stimuli that induce this type of emotions) would be processed mainly by the left hemisphere, while negative emotions would be mainly processed by the right hemisphere (Davidson et al., 1992; Canli et al., 1998). Quite similarly, Flores-Gutiérrez and collaborators (2007), in whose study participants listened to music with positive or negative emotional value, found that “unpleasant” music was mainly elaborated by the right hemisphere, while the “pleasant” music mainly activated the right hemisphere (Flores-Gutiérrez et al., 2007).

The right lateralized temporal activation for stimuli with negative valence was also found for negative affective vocalizations (Proverbio et al., 2020a) and speech utterances (Proverbio et al., 2020b) as well as in the fMRI study by Belin and collaborators (2008). In this latest study, Belin and colleagues analyzed the brain responses of individuals who listened to animal sounds (monkeys and cats) recorded in positive contexts or in negative contexts. It was found that, although participants were not able to explicitly distinguish between the “positive” and “negative” affective vocalizations of animals, the two types of stimuli were clearly distinguished in their brain (Belin et al., 2008). Through fMRI, they discovered that there was a difference in processing at the level of the secondary auditory cortex: in both hemispheres, it was found a greater activation for negative (animal or human) vocalizations compared to positive ones. This could depend on the fact that there are important structural differences between stimuli with different affective values: in general, negative vocalizations are characterized by longer sounds that activate the auditory cortex more effectively. The prefrontal cortex showed an opposite response pattern: the bilateral pars orbitalis (area 47 of Brodmann, inferior frontal gyrus) was more activated by positive vocalizations of animals. This could reflect the elaboration of the acoustic differences associated with valence, at a higher level of integration than the auditory cortex. Another region engaged in the processing of valence was the orbitofrontal cortex (OFC), with right OFC responding more to negative (human or vocal) than positive vocalizations (Belin et al., 2008).

The present data allowed us to conclude that there are specific neural mechanisms subserving the recognition of the emotional tone of music, and that they are partly shared with those devoted to understanding speech prosody and affective vocalizations. In particular, a significant difference in stimulus processing was identified starting from 600-700 ms, between 1100 and 1200 ms and between 1700 and 1800 ms. A lateralization effect was found for the activation of the temporal cortex, also shown by previous neuroimaging studies, with the medial and inferior portion of the left temporal lobe being activated more during the processing of melodies with a positive connotation. On the contrary, the right temporal lobe, in particular the rSTS (BA42), was more activated during the processing of melodies with a negative affective valence. This pattern of activation fits with previous knowledge on the neural circuits involved in the comprehension of affective prosody. A hierarchical model has been proposed by various authors (Ethofer et al., 2006; Bach et al., 2008; Lattner et al., 2005; Brück et al., 2011; Zhang et al.,2018), according to which the first stage of processing, the extraction of acoustic parameters, would involve the auditory cortex (the medial and the superior temporal cortex). Secondly, the posterior part of the right superior temporal cortex would identify the affective nature of prosody by means of multimodal integration. Finally, the bilateral inferior frontal gyrus and the orbitofrontal cortex would perform a sophisticated evaluation and semantic understanding of the vocally expressed emotions. It can be hypothesized that broadly the same circuits would be stimulated by the melodic variations of musical information.

Indeed, the results of the present study hints at the existence of a common circuit for processing the prosodic aspect of language and music, with different activations based on the affective valence of the stimuli, regardless of their timbre. Figure 5 shows a plot of common activations found in the present as well as in other ERP and neuroimaging studies contrasting auditory stimuli as a function of their emotional valence.

**Figure 5.**
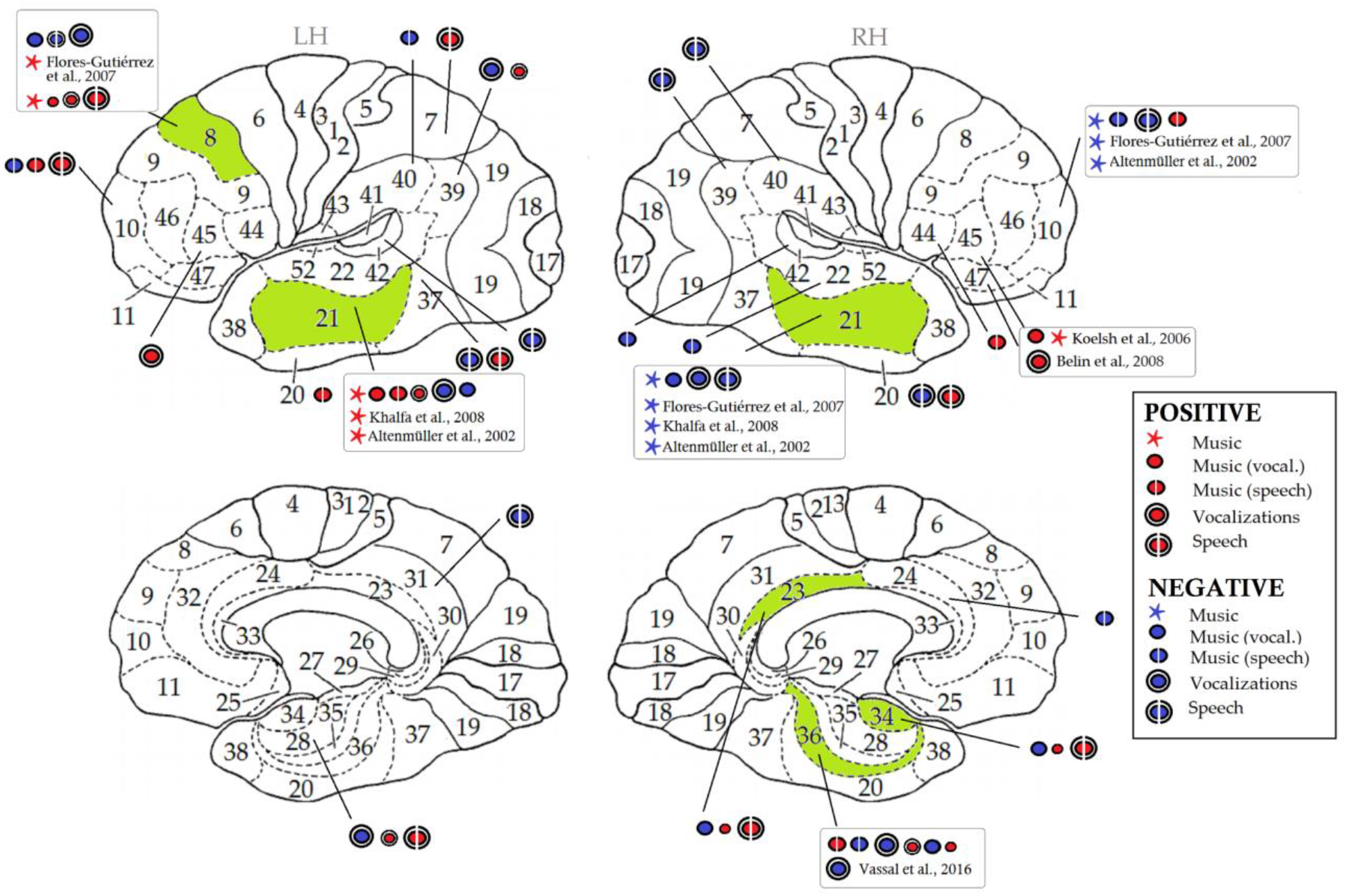
Schematic representation of brain activations associated with listening to positive and negative music, affective vocalizations, and speech according to recent electrical neuroimaging and fMRI literature. The shaded green areas indicate common activations for musical and affective aspects of language in the present and in other two ERP studies exploring the neural bases of comprehension of: vocalizations, music from vocalizations (Proverbio et al., 2020a), speech (Proverbio et al., 2020b) and music from speech (present data). Therefore, an area activated uniquely by speech (e.g. the left BA42) will not appear as shaded in this picture. Koelsch et al., (2006) found an activation of the inferior frontal gyrus during listening to positive music. In Belin and coauthors’ paper (2008) BA47 was more active during listening to positive vocalizations of animals. The right OFC responded more to negative vocalizations than positive ones and this happened independently of the source of origin, humans or animals (Belin et al., 2008). In Flores-Gutiérrez et al.’s fMRI study (2007) pleasant music activated the left medial prefrontal area and the left posterior temporal-parietal region, while unpleasant music activated the right superior frontopolar region, the insula (slightly asymmetrically) and the right auditory area. Khalfa et al. (2008) examined patients with right or left anterior medial temporal resections and found that patients with left lesions showed a poorer ability to identify happy music, while patients with resection of the right anteromedial temporal lobe showed deficits in the recognition of sadness. In Altenmüller et al.’s study (2002) it was found that positive emotional content of music was accompanied by an increase in left temporal activation, while a negative content was associated with a preponderance of the right fronto-temporal cortex activation. In Vassal et al.’s study (2016) the right uncus (BA 36) was associated with listening to vocalizations. Red circles refer to positive emotions, blue circles to negative emotions.

The proposed model might contribute to understand while there are so many universalities in the aesthetical and emotional effects of music listening (for example, the simple happy vs. sad distinction: Bogert et al., 2016; Curtis et al., 2010; Mitterschiffthaler et al., 2007), regardless of culture, familiarity and style. Quite interestingly, it has been showed (Sachs et al., 2016) that white matter connectivity between the superior temporal gyrus, insula and medial prefrontal cortex explains individual differences in reward sensitivity to music. This universality has to do with an innate mechanism for comprehending prosody and language in the humans species, according to an evolutionary neurobiological perspective (Panksepp and Bernatzky, 2002; Cook et al., 2006).

## ACKNOWLEDGEMENTS

This study was funded by 2017-ATE-0058 grant n° 16940 by University of Milano-Bicocca to AMP, entitled “The processing of emotional valence in language and music: an ERP investigation”. We are extremely grateful to sound engineer Roberto Oldani of *Claudio Abbado Civic School of Music* of Milan for transforming the stimuli in music. The authors are deeply grateful to Francesco de Benedetto and Sacha Santoni for their technical support.

